# Brain-Specific Inhibition of mTORC1 by a Dual Drug Strategy: A Novel Approach for the Treatment of Alcohol Use Disorder

**DOI:** 10.1101/2020.10.12.336701

**Authors:** Yann Ehinger, Ziyang Zhang, Khanhky Phamluong, Drishti Soneja, Kevan M. Shokat, Dorit Ron

**Author notes:** YE and ZZ contributed equally to the manuscript. Corresponding author: (DR).

## Abstract

Alcohol Use Disorder (AUD) is a devastating psychiatric disorder affecting a large portion of the population. Unfortunately, efficacious medications to treat the disease are limited and thus AUD represents an area of unmet medical need. mTORC1 plays a crucial role in neuroadaptations underlying alcohol use. mTORC1 also contributes to alcohol craving, habit, and relapse. Thus, mTORC1 inhibitors are promising therapeutic agents to treat AUD. However, chronic inhibition of mTORC1 in the periphery produces undesirable side effects in humans, which limit their potential clinical use for the treatment of AUD. To overcome these limitations, we utilized a binary drug strategy in which mice were co-administered the mTORC1 inhibitor RapaLink-1 together with a novel small molecule (RapaBlock) to protect mTORC1 activity in the periphery. We show that the dual administration of RapaLink-1 with RapaBlock, abolishes RapaLink-1-dependent mTORC1 inhibition in the liver and blocks adverse side effects detected in humans including body weight loss, glucose intolerance and liver toxicity. Importantly, we show that co-administration of RapaLink-1 and RapaBlock inhibits alcohol-dependent mTORC1 activation in the Nucleus Accumbens and robustly moderates the level of alcohol use. Our data present a novel approach that could be used to treat individuals suffering from AUD.

---

Alcohol Use Disorder (AUD) is characterized by compulsive alcohol intake despite negative consequence^1,2^. AUD is widespread; afflicting 10-15% of the population and causing significant medical, social, and economic burdens^2–4^. In fact, AUD is one of the most prevalent mental health disorders^2^. Furthermore, the incidence of AUD diagnosis has increased by 35% in the United States between 2001 and 2013^1^. Unfortunately, pharmacotherapeutic options for treating AUD are rather limited^5^. Only 3 drugs, naltrexone, acamprosate and disulfiram, have been approved by the US Food and Drug Administration (FDA) for AUD treatment, unfortunately however, these medications suffer from efficacy issues and produce detrimental side effects^2^. Thus, there is an urgent need to develop effective medications to treat phenotypes such as binge drinking, craving, and relapse.

Mechanistic target of Rapamycin complex 1 (mTORC1) represents an ideal drug target for the treatment of AUD. mTORC1 is a multiprotein complex that contains the serine/threonine protein kinase mTOR and adaptor proteins, including Raptor, Deptor and mLST8^6,7^. mTORC1 is activated by growth factors, amino acids, and oxygen^7,8^, and plays a role in lipid genesis, glucose homeostasis, protein translation^7,8^ and autophagy^9^. Hyperactivation of mTORC1 has been linked to pathological states such as insulin resistance and cancer^7,8^. In the CNS, mTORC1 is activated by neurotransmitters and neuromodulators, such as glutamate and BDNF^6,10^. Upon activation, mTORC1 phosphorylates eIF4E-binding protein (4E-BP) and the ribosomal protein S6 kinase (S6K), which in turn phosphorylates its substrate, S6^7^. These phosphorylation events precede the initiation of local dendritic translation of synaptic proteins^11,12^. As such, mTORC1 plays an important role in synaptic plasticity, and learning and memory^6,13^. mTORC1 malfunction in the CNS has been linked to aging processes^14^, neurodegenerative diseases such as Alzheimer’s disease, Parkinson’s disease and Huntington’s disease^8,13,15^, neurodevelopmental disorders such as autism, as well as psychiatric disorders including addiction^13,14,16^. Growing evidence implicates mTORC1 in mechanisms underlying AUD^17^. Specifically, using Rapamycin, a selective allosteric mTORC1 inhibitor^18^, and RapaLink-1, a third generation mTORC1 inhibitor^19^, in combination with rodent paradigms that model phenotypes associated with AUD, mTORC1 was found to contribute to mechanisms underlying alcohol seeking, excessive alcohol consumption^20,21,22^, reward memory, and habitual alcohol use^10,20,21^. Finally, mTORC1 also plays a crucial role in mechanisms that drive relapse to alcohol drinking^23,24^, a crucial step in the addiction cycle^25^.

Because of the important role of mTORC1 in various pathological states, the kinase represents an attractive drug target for the treatment of numerous diseases. Indeed, Rapamycin and its analogs (Rapalogs) have been approved by the FDA for the prevention of host rejection after transplantation^26,27^, as well as for the treatment of for several types of cancer, Tuberous Sclerosis, and cardiovascular disease^28,29^. However, chronic inhibition of mTORC1 in the periphery produces detrimental side effects such as suppression thrombocytopenia, impaired glucose sensitivity, hyperlipodemia, decreased wound healing and the suppression of the immune system^30 28^, thus limiting the utility of Rapamycin and other Rapalogs for the treatment of CNS disorders such as AUD because of safety concerns.

In an attempt to circumvent these undesirable effects resulting from sustained mTORC1 inhibition in the periphery, we developed an approach which enables CNS-specific inhibition of mTORC1 while protecting the activity of the kinase in the periphery (**Fig. 1**)^31^. Specifically, we utilized the unique mechanism of action of the mTORC1 inhibitors, Rapamycin and RapaLink-1, which requires their binding to the chaperone, FK506 binding protein 12 (FKBP12) prior to the inhibition of the kinase (**Fig. 1**)^32^. We designed a brain impermeable small molecule (Rapablock) that binds FKBP12, and acts to prevent access to the necessary factor for mTOR inhibition (**Fig. 1**). We hypothesized that when Rapablock will be co-administered with RapaLink-1, mTORC1 activity will be protected in the periphery while inhibited in the brain (**Fig. 1**)^31^. We further predicted that this approach will block the undesirable side effects observed after chronic inhibition of the kinase. Finally, we tested the utility of the approach in a preclinical mouse model of AUD.

**Fig. 1.**
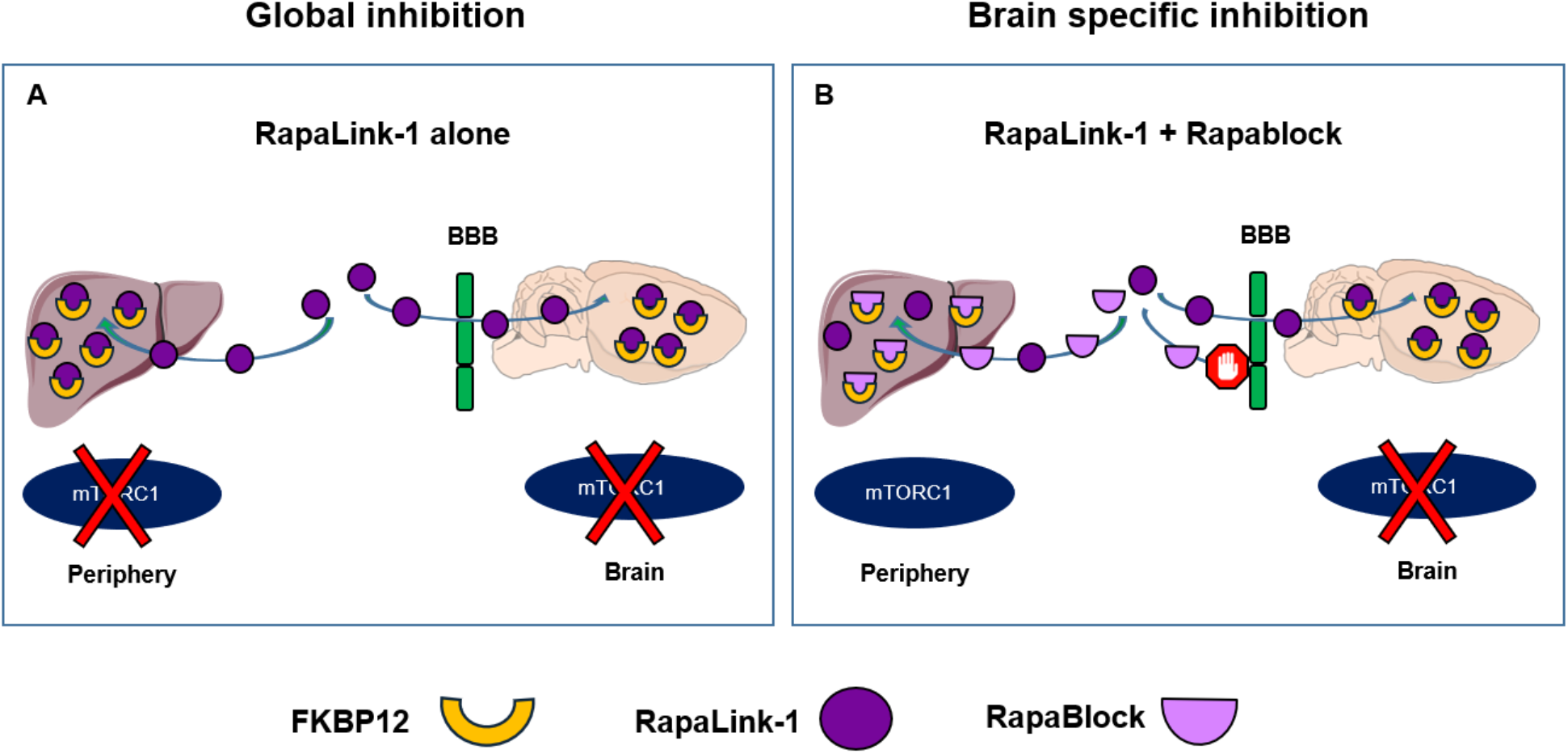
Schematic representation of strategy. **(A)** Systemic administration of RapaLink-1 (purple) inhibits mTORC1 in the periphery and in the brain. **(B)** RapaBlock (pink), a small molecule that does not cross the blood brain barrier (BBB) (green) and competes with RapaLink-1 (purple) for FKBP12 (yellow) binding in the periphery thereby selectively protects mTORC1 activity outside of the CNS. Systemic co-administration of RapaLink-1 and RapaBlock allows brain specific inhibition of mTORC1.

First, to determine if RapaBlock protects mTORC1 activity in the periphery, mice received a systemic administration of RapaLink-1 alone (1mg/kg) or a combination of RapaLink-1 and RapaBlock (40mg/kg), and mTORC1 activity in the periphery was measured 3 hours later (**Fig. 2A**). The liver was chosen as a peripheral organ since the mTORC1 pathway plays a critical role in hepatic function^7^, and since chronic inhibition of mTOR in the liver has been implicated in liver toxicity^33^. As expected, RapaLink-1 administrated alone blocked the phosphorylation of the mTORC1 downstream targets, S6 (**Fig. 2B,C**) and 4E-BP (**Fig. 2D,E**) in the liver. In contrast, co-administration of RapaLink-1 and RapaBlock produced a complete protection of mTORC1 activity in the liver (**Fig. 2B-E**) demonstrating that RapaBlock protects mTORC1 activity in the periphery.

**Fig. 2.**
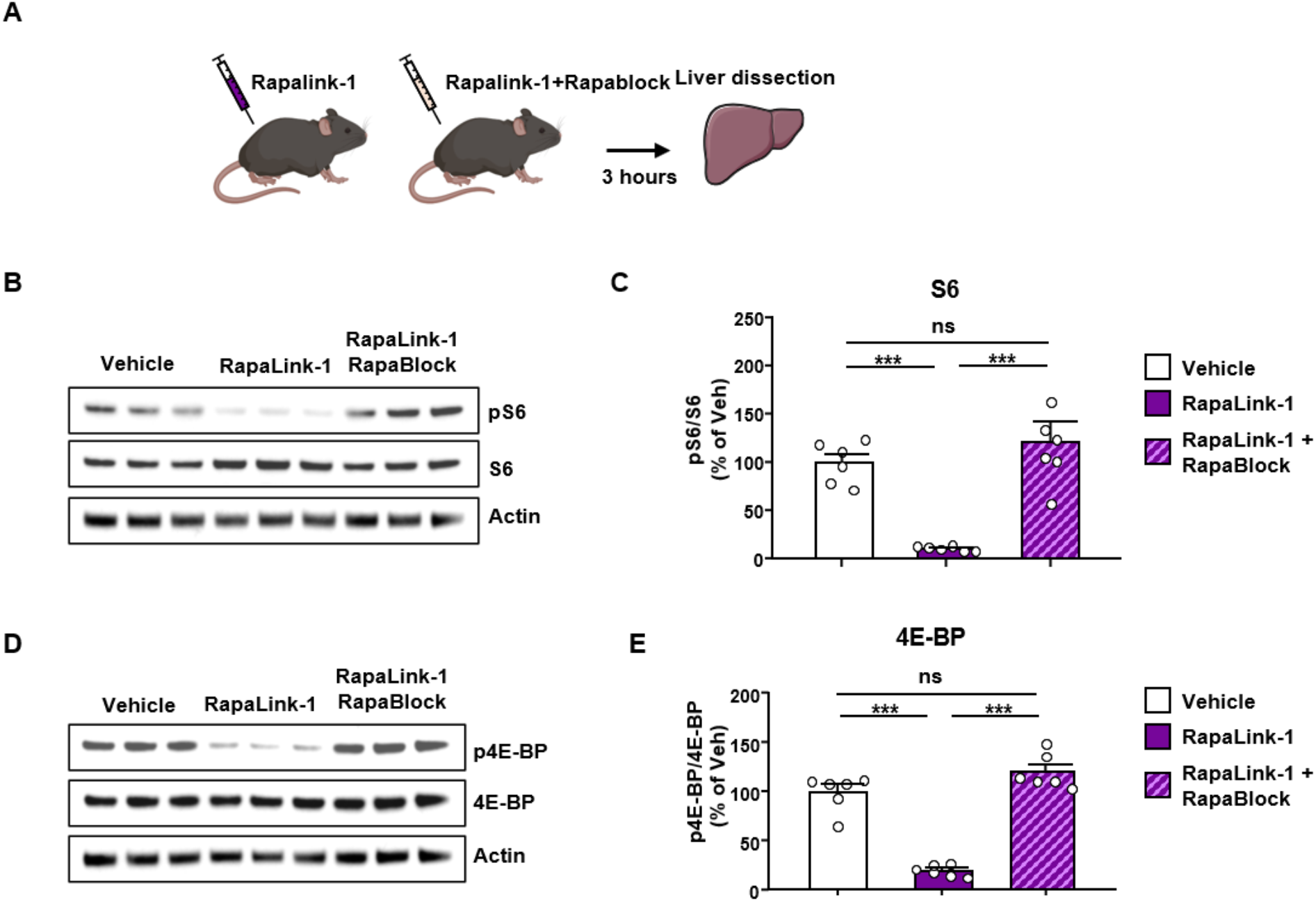
RapaBlock protects mTORC1 activity in the liver. **(A)** Timeline of experiment. Mice received a systemic administration of RapaLink-1 alone (1 mg/kg) or a combination of RapaLink-1 (1 mg/kg, purple) and RapaBlock (40 mg/kg, pink), and mTORC1 activity in the liver was measured 3 hours later. **(B, D)** Representative images depict S6 phosphorylation (pS6) (**B**) and 4E-BP phosphorylation (p4E-BP) (**D**) (top panels), total protein levels of S6 (**B**) and 4E-BP (**D**) (Middle panels), and actin (B,**D**) (bottom panels), which was used as a loading control. **(C,E)** Data are presented as the individual data points and mean densitometry values of the phosphorylated protein divided by the densitometry values of the total protein ± SEM and expressed as % of vehicle. Significance was determined using One-way ANOVA followed by Tukey’s multiple comparisons test. Co-administration of RapaLink-1 and RapaBlock protects S6 (One-way ANOVA: *F*_2,15_ = 20.56, *P* < 0.0001) and 4E-BP (One-way ANOVA: *F*_2,15_ = 42.77, *P* < 0.0001) phosphorylation in the liver. n=6 per condition. ***p<0.001, ns = non-significant.

As mentioned above, chronic administration of Rapamycin produces a broad range of undesirable effects such as body weight loss, impaired glucose metabolism^28,34^ and liver toxicity^33^. We next examined whether RapaBlock could protect against these adverse effects caused by chronic mTORC1 inhibition in the periphery. Mice were chronically treated 3 times a week for 4 weeks with either RapaLink-1 alone (1mg/kg) or with a combination of RapaLink-1 (1mg/kg) and RapaBlock (40mg/kg) (**Fig. 3A**). Similar to what was previously reported for Rapamycin^34^, chronic treatment of mice with RapaLink-1 led to a significant decrease in body weight (**Fig. 3B)**. However, the combination of RapaLink-1 and RapaBlock prevented the decrease in the weight of the mice (**Fig. 3B)**.

**Fig. 3.**
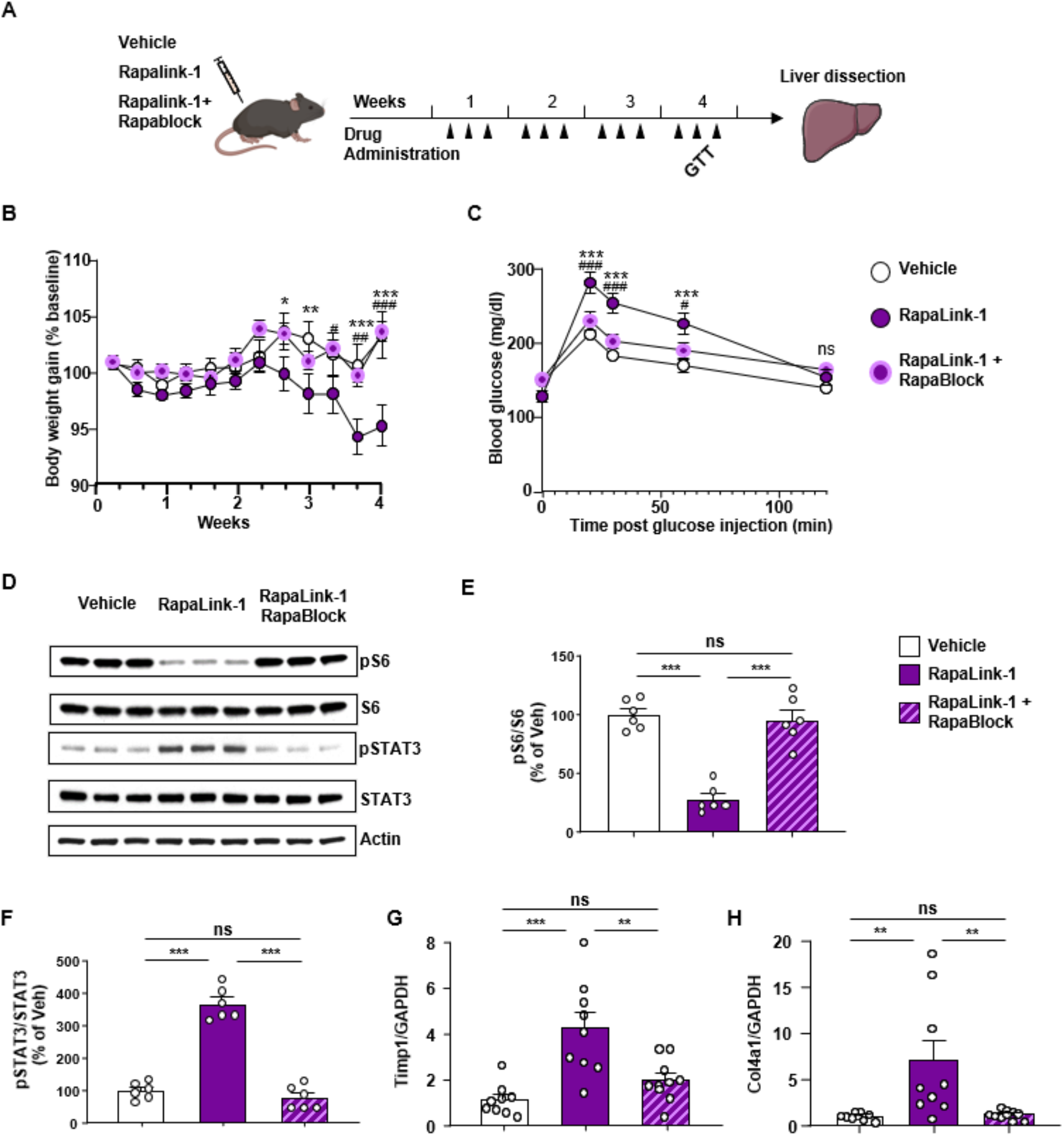
RapaBlock abolishes RapaLink-1-dependent weight loss, glucose intolerance and liver toxicity. **(A)** Timeline of experiment. Mice were treated 3 times a week with RapaLink-1 alone (1 mg/kg) (purple) or with a combination of RapaLink-1 (1 mg/kg, purple) and RapaBlock (40 mg/kg, (pink) for 4 weeks, and body weight, glucose tolerance and liver toxicity were evaluated. **(B)** Co-administration of RapaLink-1 and RapaBlock eliminates RapaLink-1-dependent body loss (Two-way ANOVA, effect of time (*F*_4,130_ = 52.13, *P* < 0.0001), effect of treatment (*F*_4,130_ = 52.13, *P* < 0.0001) and interaction (*F*_8,130_ = 5.057, *P* < 0.0001)). **(C)** Glucose tolerance test was performed during the last week of chronic drug treatment. Co-administration of RapaLink-1 and RapaBlock reduces RapaLink-1-dependent increase in blood glucose (Two-way ANOVA, effect of time (*F*_11,312_ = 3.467, *P* = 0.0001), effect of treatment (*F*_2,312_ = 27.55, *P* < 0.0001) and interaction (*F*_22,312_ = 1.731, *P* = 0.0233)). (**D-H**) Co-administration of RapaLink-1 and RapaBlock protects against RapaLink-1-dependent liver toxicity. (**D-F**) The liver was dissected 24 hours after the last drug administration and S6 and STAT3 phosphorylation were measured. **(D)** Representative images of pS6, total S6 (top panels), phospho-STAT3 (pSTAT), total STAT3 (middle panels) and actin (bottom panel). **(E)** Co-administration of RapaLink-1 and RapaBlock protects mTORC1 activity in the liver (One-way ANOVA: *F*_2,15_=42.77, *P*<0.0001). **(F)** RapaBlock reversed RapaLink-1-dependent increase in STAT3 phosphorylation (One-way ANOVA: *F*_2,15_=104.9, *P*<0.0001). **(G,H)** RapaBlock protects against RapaLink-1-dependent increase of fibrogenic markers, Timp1 (One-way ANOVA: *F*_2,25_=13.18, *P*=0.0001), and Col4a1 (One-way ANOVA: *F*_2,25_=7.446, *P*=0.0029). **(B-C)** Data are presented as mean ± SEM. Significance was determined using RM Two-way ANOVA followed by Tukey’s multiple comparisons test. Vehicle, n=9, RapaLink-1, n=10; RapaLink-1 + RapaBlock, n=10, * = RapaLink-1 vs. Vehicle, # = RapaLink-1 vs. RapaLink-1+RapaLink. * or # p<0.05, ** or ## p<0.01 and *** or ### p<0.001, ns = non-significant. **(E-F)** Data are presented as the individual data points and mean densitometry values of the phosphorylated protein divided by the densitometry values of the total protein ± SEM and expressed as % of vehicle. Significance was determined using One-way ANOVA followed by Tukey’s multiple comparisons test. n=6 per condition, ***p<0.001, ns = non-significant. **(G,H)** Data are expressed as a ratio to total GAPDH and presented as individual data points and mean ± SEM. Significance was determined using One-way ANOVA followed by Tukey’s multiple comparisons test. Vehicle, n=9, RapaLink-1, n=9; RapaLink-1 + RapaBlock, n=10, **p<0.01, ***p<0.001, ns = non-significant.

As chronic inhibition of mTORC1 in the periphery has been linked to hyperglycemia and insulin resistance^34,35^, we determined whether long-term administration of RapaLink-1 causes glucose intolerance and whether RapaBlock blocks this effect. To do so, a fasting glucose tolerance test (GTT) was conducted during the fourth week of treatment of mice with RapaLink-1 (1mg/kg) alone or RapaLink-1 (1mg/kg) and RapaBlock (40mg/kg). Blood glucose levels were markedly increased in mice chronically treated with RapaLink-1 (**Fig. 3C**). In contrast, blood glucose levels were similar in vehicle-treated vs. RapaLink-1/RapaBlock-treated mice suggesting the RapaBlock also protects against the glucose intolerance side effects.

Prolonged treatment with Rapamycin in mice results in liver inflammation^33^. To determine whether chronic administration of RapaLink-1 produces a similar liver toxicity phenotype, the liver was dissected and harvested following four weeks of RapaLink-1 treatment, and liver inflammation was evaluated by measuring the phosphorylation level of the signal transducer and activator of transcription 3 (STAT3), a prerequisite for the activation of the transcription factor, as well as the expression of the fibrogenic markers, tissue inhibitor of metalloproteinase 1 (Timp1), and collagen α1(IV) (Col4A1)^33^. We found that chronic administration of RapaLink-1 while inhibiting mTORC1 activity (**Fig. 3D,E**), robustly elevated the level of STAT3 phosphorylation at the Tyrosine 705 site (**Fig. 3D,F**), Four weeks administration of RapaLink-1 also increased the mRNA levels of *Timp1* (**Fig. 3G**) and *Col4A1* (**Fig. 3H**), suggesting that the drug produces liver toxicity. Importantly, RapaBlock protected mTORC1 activity in the liver (**Fig. 3D,E**) and at the same time, RapaBlock prevented the increase in STAT3 phosphorylation (**Fig. 3D,F**), and the expression of liver toxicity markers (**Fig. 3G-H**), suggesting that RapaBlock eliminates liver toxicity issues associated with prolonged inhibition of mTORC1.

Together, our results suggest that similar to Rapamycin^28^, RapaLink-1 on its own has limited utility due to significant adverse side effects such as reduction in body weight, glucose intolerance and liver toxicity. However, the finding that RapaBlock is able to fully prevent adverse effects resulting from the sustained inhibition of mTORC1 in the periphery suggests that this approach could be beneficial for patients.

To examine the utility of the approach for a CNS application, we first examined the behavioral consequences of chronic systemic administration of RapaBlock alone. We reasoned that if RapaBlock does not cross the blood brain barrier (BBB) it should produce no adverse cognitive effects when administrated chronically to mice. We therefore treated mice with RapaBlock (40mg/kg) for 6 weeks and performed a battery of behavioral tests following 3 and 6 weeks of treatment (**Fig. S1A**). Chronic RapaBlock treatment did not alter sensorimotor coordination, as measured by the latency to fall from a rotarod apparatus (**Fig. S1B-D**), anxiety-like behavior, as measured in an elevated plus maze paradigm (**Fig. S1E-G**), or recognition memory, which was tested using a novel object recognition paradigm (**Fig. S1H-I**). These results suggest that RapaBlock has no behavioral effects on its own.

Alcohol activates mTORC1 in numerous brain regions including the nucleus accumbens (NAc)^20,36^, and blockade of mTORC1 in the CNS attenuates numerous phenotypes associated with alcohol use including excessive alcohol intake^10,17,21,24^. Having demonstrated the ability of RapaBlock to protect the function of mTORC1 in the periphery and to block the adverse phenotypes stemming from chronic RapaLink-1 treatment without producing behavioral side effects, we examined whether this approach can produce a selective inhibition of mTORC1 activity in the brain and be a potential effective treatment for AUD. To do so, we first determined whether RapaLink-1 inhibits alcohol-dependent mTORC1 activity in the NAc. Mice underwent 7 weeks of intermittent access to 20% alcohol in a two-bottle choice, a paradigm that models binge alcohol intake in humans^37^. Mice consuming water only for the same length of time were used as a control. On week 8, mice received a systemic administration of RapaLink-1 alone (1mg/kg) or a combination of RapaLink-1 (1mg/kg) and RapaBlock (40mg/kg) 3 hours before the last 24-hour drinking session, at the conclusion of which mTORC1 activity was measured in the NAc (**Fig. 4A**). Excessive alcohol intake increased phosphorylation of the mTORC1 targets S6 and 4-EBP in the NAc of mice consuming alcohol and treated with vehicle vs. animals consuming water only (**Fig. 4B-E**). Systemic administration of RapaLink-1 inhibited alcohol-dependent mTORC1 activation in the NAc in the absence or presence of RapaBlock (**Fig. 4B-E**). In contrast, systemic administration of RapaBlock alone had no effect on alcohol-dependent activation of mTORC1 in the NAc (**Fig. S2**).

**Fig. 4.**
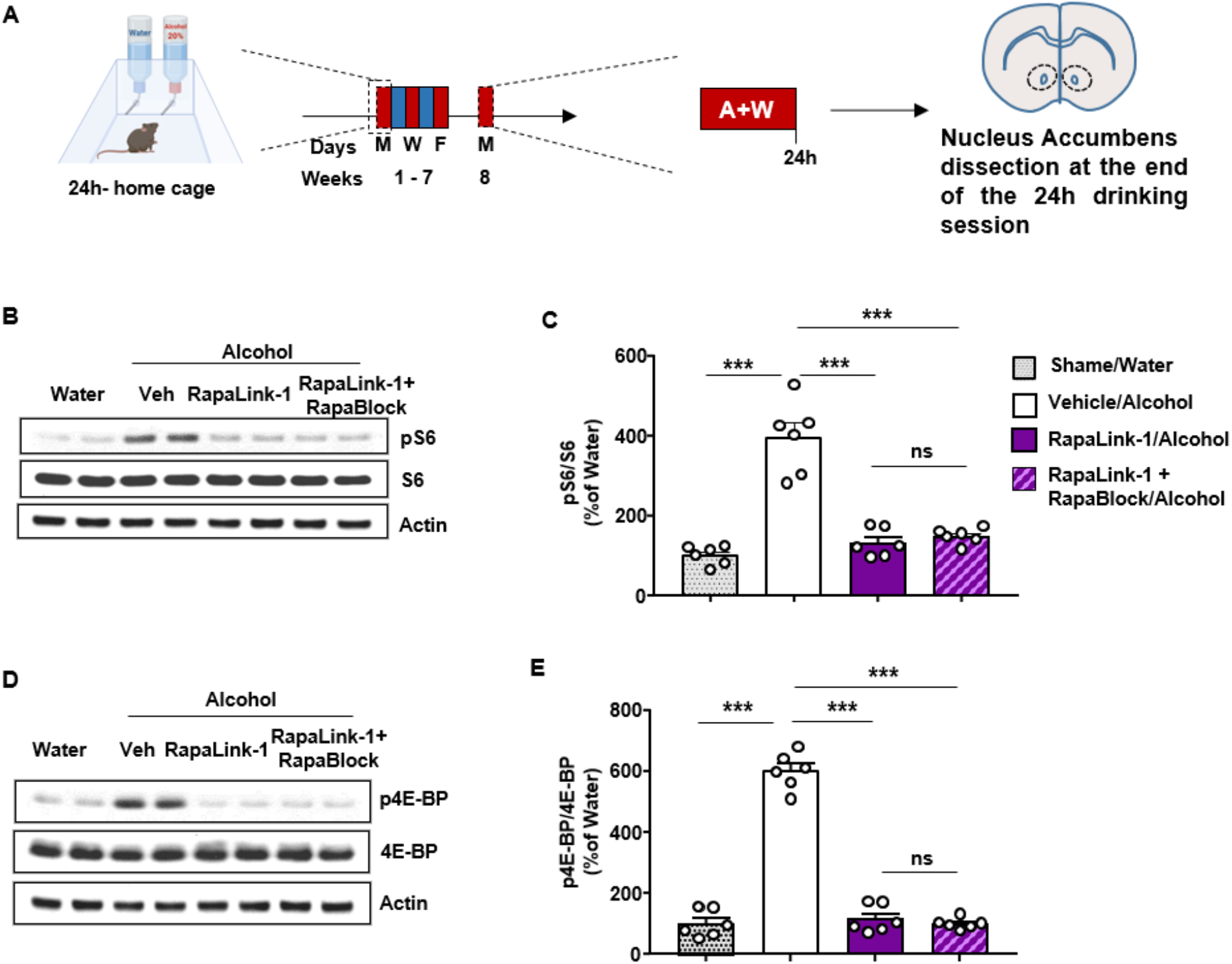
RapaLink-1 inhibits mTORC1 activity in the Nucleus Accumbens in the presence and absence of RapaBlock. **(A)** Timeline of experiment. Mice underwent 7 weeks of IA20%2BC. On week 8, mice received a systemic administration of RapaLink-1 alone (1 mg/kg, purple) or a combination of RapaLink-1 (1 mg/kg, purple) and RapaBlock (40 mg/kg, pink) 3 hours before the beginning the drinking period, and the NAc was removed at the end of the last 24 hours drinking session. **(B,D)** Representative images depict S6 phosphorylation (pS6) **(B)** and 4E-BP phosphorylation (p4E-BP) **(D)** (top panels), total protein levels of S6 **(D)**, and 4E-BP **(D)** (Middle panels) and actin (bottom panels). **(C,E)** RapaLink-1 inhibits NAc mTORC1 activity in the presence of RapaBlock, as shown by the significantly decreased levels of pS6 (One-way ANOVA: *F*_3,20_= 41.58, *P*<0.0001) and p4E-BP (One-way ANOVA: *F*_3,20_ = 186, *P*<0.0001). Data are presented as the individual data points and mean densitometry values of the phosphorylated protein divided by the densitometry values of the total protein ± SEM and expressed as % of vehicle. Significance was determined using One-way ANOVA Tukey’s multiple comparisons test. n=6 per condition. ***p<0.001, ns = non-significant.

Finally, we tested if RapaLink-1 attenuates alcohol intake in the presence or absence of RapaBlock (**Fig. 5A**). RapaBlock administered alone had no effect on alcohol and water intake (**Fig. S3**) whereas similar to what we previously reported^21^, RapaLink-1 reduced alcohol intake and reward measured during the first 4 hours of an alcohol binge drinking session (**Fig. 5B,C**), and at the end of a 24-hour session (**Fig. 5E,F**), Importantly, co-administration of RapaLink-1 and RapaBlock produced similar attenuation of alcohol intake and preference, compared to the vehicle group (**Fig. 5B,C, E,F**). The co-treatment of RapaLink-1 and RapaBlock did not affect water intake (**Fig. 5D,G**) nor did the regimen alter blood alcohol concentration (**Fig. S4**). Together, these data suggest that RapaLink-1 preserves its desirable inhibitory actions on mTORC1 in the brain and inhibits heavy alcohol use when administered together with RapaBlock.

**Fig. 5.**
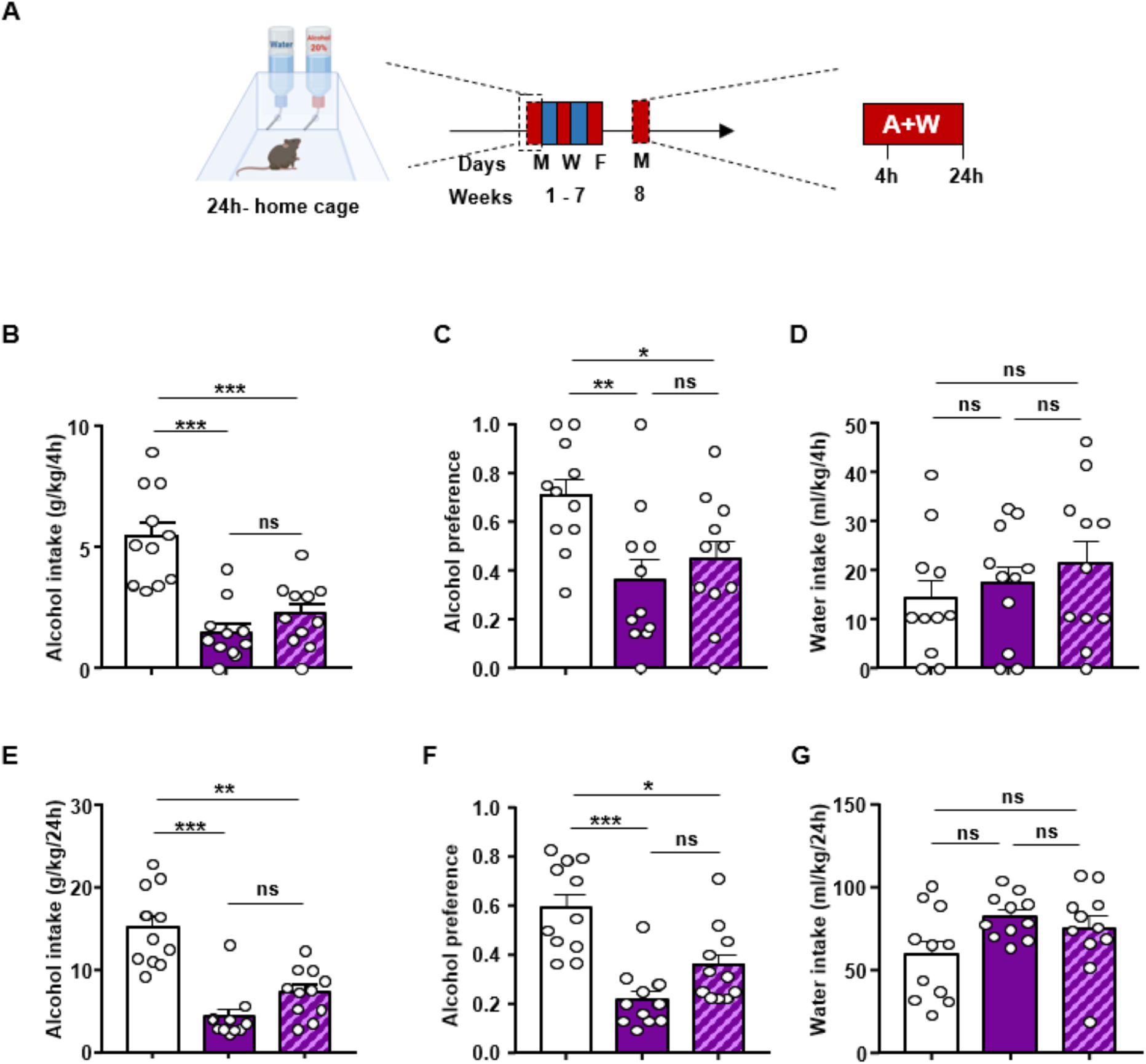
RapaLink-1 reduces alcohol intake and preference in the presence and absence of RapaBlock. **(A)** Timeline of experiment. Mice underwent 7 weeks of IA20%2BC. On week 8, mice received a systemic administration of RapaLink-1 alone (1 mg/kg, purple) or a combination of RapaLink-1(1 mg/kg, purple) and RapaBlock (40 mg/kg, pink) 3 hours before the beginning of drinking session. Alcohol and water intake were measured 4 hours **(B,D)** and 24 hours later **(E,G)**. Alcohol preference was calculated as the ratio of alcohol intake to total fluid intake at the end of the 4-hour (**C)** and 24-hour (**F)** drinking session. **(B-C)** Administration of RapaLink-1 alone or RapaLink-1+RapaBlock significantly decreased alcohol intake (One-way ANOVA: *F*2,30 = 20.46, *P* < 0.0001) and alcohol preference (One-way ANOVA: *F*_2,30_ = 5.482, *P* = 0.0094) at the end of a 4-hour session (**D**) Water intake at the end of a 4-hour session was not affected by treatments (One-way ANOVA: *F*_2,30_ = 0.7708, *P* = 0.4716). (**E-F**) Administration of RapaLink-1 alone or RapaLink-1+RapaBlock significantly decreased alcohol intake (One-way ANOVA: *F*_2,30_ = 25.47, *P* < 0.0001) and alcohol preference (One-way ANOVA: *F*_2,30_ = 16.73, *P* < 0.0001) at the end of a 24-hour session. (**G**) Water intake at the end of a 24-hour session was not affected by treatments (One-way ANOVA: *F*_2,30_ = 2.966, *P* = 0.0668). Data are presented as individual data points and mean ± SEM. Significance was determined using One-way ANOVA Tukey’s multiple comparisons test. n=11 per condition. *p<0.05, **p<0.01, ***p<0.001 and ns = non-significant.

## Conclusions

We show herein that RapaBlock provides full protection of mTORC1 activity in the periphery, and by so doing prevents the detrimental side effects resulting from chronic inhibition of the kinase in the periphery. We further provide proof of concept data for the potential utility of the RapaLink-1/RapaBlock dual drug administration strategy for the treatment of AUD.

Our data suggest that RapaBlock while acting in the periphery, the drug is inert in the CNS. Specifically, RapaBlock did not affect mTORC1 activity in the brain, neither did it alter alcohol and water intake or affected mice’s locomotion, learning and memory, anxiety-like behavior. Importantly, RapaBlock prevented numerous adverse side effects resulting from long-term inhibition of mTORC1 in the periphery. However, more studies are warranted to test RapaBlock’s ability to prevent other major side effects produced in humans by mTORC1 inhibitors such as immunosuppression and diabetes^38^. Nevertheless, our data suggest that RapaBlock in combination with RapaLink-1 could be used for CNS-specific applications such as AUD.

Furthermore, the approach described herein, enabling the separation between the desirable, CNS-mediated actions of a drug versus the undesirable periphery-mediated drug effects could be used for the development of other CNS-targeted therapeutic approaches. For instance, Fyn kinase has been implicated in mechanisms underlying Alzheimer’s disease^39^, AUD^40^ and opiate addiction^41^, and small molecule inhibitors such as AZD0530 have been in development for the treatment of Alzheimer’s disease^39^. Protecting the activity of the kinase in the periphery will enable the reduction of potential side effects and increase the safety of the inhibitor.

AUD is the third most preventable disease^3^, unfortunately, drug development for the treatment of AUD has only been modestly successful to date^2^. Data obtained in rodents suggest that inhibition of mTORC1 in the brain dampens numerous adverse behaviors associated with alcohol use^10,17,21,24^. In contrast, inhibition of mTORC1 does not alter the consumption of natural rewarding substances^40^, suggesting that inhibition of mTORC1 does not affect reward *per se*. Furthermore, inhibition of mTORC1 is not aversive or rewarding, nor does it alter locomotion^40^. Importantly, a single administration of Rapamycin or RapaLink-1 produces a long-lasting attenuation of excessive alcohol use and relapse^21,23^. Putting together these findings with the data presented herein strongly suggests that Rapalog/Rapablock dual drug strategy should be tested in humans.

Interestingly, mTORC1 has been linked to neuroadaptations associated with numerous drugs of abuse^16^. For instance, Rapamycin administration was shown to inhibit reconsolidation of cocaine and morphine reward memory as well as the reinstatement of cocaine self-administration^16^. In addition, treatment of rodents with Rapamycin was reported to inhibit consolidation and reconsolidation of fear memory^42,43^. Together, these findings raise an attractive possibility that the Rapalog/RapaBlock dual drug strategy could be developed as a therapeutic option not only for AUD but also for the treatment of addiction to other drugs of abuse as well as for post-traumatic stress disorder.

## Acknowledgments

The authors thank Dr. Jeffrey Moffat for his input. This research was supported by the National Institute of Alcohol Abuse and Alcoholism, R01 AA027474 (DR). K.M.S. acknowledges NIH 1R01CA221969, the Michael J. Fox Foundation P0536220, The Samuel Waxman Research Foundation and the Howard Hughes Medical Institute for support.

(KMS and ZZ) are co-inventors on patent applications covering RapaBlock owned by UCSF. KMS is an inventor on patents covering Rapa-Link1 owned by UCSF and licensed to Revolution Medicines. KMS receives monetary and stock compensation and is a co-founder and SAB member of Revolution Medicines.

The other authors do not have financial or non-financial competing interests.

## Materials and Methods

### Animals

Male C57BL/6J mice (Jackson Laboratory, Bar Harbor, ME) were 6-7 weeks old at the beginning of the experiment and were individually housed in temperature- and humidity-controlled rooms under a reversed 12-hour light/dark cycle (lights on at 10:00 PM), with food and water available ad libitum. All animal procedures in this report were approved by the University of California San Francisco (UCSF) Institutional Animal Care and Use Committee and conducted in agreement with the Association for Assessment and Accreditation of Laboratory Animal Care (AAALAC, UCSF).

### Reagents

Anti-phospho-S6 (S235/236, 1:500), anti-S6 (1:1000), anti-phospho-4E-BP (T37/46, 1:500), anti-4E-BP (1:1000), anti-phosphoTyr705-STAT3 (1:500) and anti-STAT3 (1:500) antibodies were purchased from Cell Signaling Technology (Danvers, MA). Anti-Actin (1:10,000) antibodies, phosphatase Inhibitor Cocktails 2 and 3, and Dimethyl sulfoxide (DMSO) were purchased from Sigma Aldrich (St. Louis, MO). Nitrocellulose membrane was purchased from EMD Millipore (Billerica, MA, USA). Enhanced Chemiluminescence (ECL) was purchased from GE Healthcare (Pittsburg, PA). Donkey anti-rabbit horseradish peroxidase (HRP) and donkey anti-mouse horseradish peroxidase (HRP) were purchased from Jackson ImmunoResearch (West Grove, PA). AMV reverse transcriptase was purchased from Promega (Madison, WI). SYBR Green PCR Master mix was purchased from Thermo Fisher Scientific, Inc. (Waltham, MA, USA). EDTA-free complete mini Protease Inhibitor Cocktail was purchased from Roche (Indianapolis, IN). NuPAGE Bis-Tris precast gels and Phosphate buffered saline (PBS) were purchased from Life Technologies (Grand Island, NY). Bicinchoninic Acid (BCA) protein assay kit was obtained from Thermo Scientific (Rockford, IL). ProSignal Blotting Film was purchased from Genesee Scientific (El Cajon, CA). Ethyl alcohol (190 proof) was purchased from VWR (Radnor, PA).

### Tissue harvesting

Animals were euthanized and the brain and liver were rapidly removed an anodized aluminum block on ice. The NAc was isolated from a 1 mm thick coronal section located between +1.7 mm and +0.7 mm anterior to bregma according to the Franklin and Paxinos stereotaxic atlas (3rd edition). Collected tissues were immediately homogenized in 300 ul RadioImmuno Precipitation Assay (RIPA) buffer containing (in mM: 50 Tris-HCl, pH 7.6, 150 NaCl, 2 EDTA), and 1% NP-40, 0.1% SDS and 0.5% sodium deoxycholate and protease and phosphatase inhibitor cocktails. Samples were homogenized by a sonic dismembrator. Protein content was determined using a BCA kit.

### Western Blot analysis

Equal amounts of homogenates from individual mice (30 ug) were resolved on NuPAGE Bis-Tris gels and transferred onto nitrocellulose membranes. Blots were blocked in 5% milk-PBS, 0.1% Tween 20 for 30 minutes and then incubated overnight at 4°C with anti-pS6, anti-p4E-BP and anti-p[Y705]STAT3 antibodies. Membranes were then washed and incubated with HRP-conjugated secondary antibodies for 2 hours at room temperature. Bands were visualized using ECL. Membranes were then incubated for 30 minutes at room temperature in a stripping buffer containing 25 mM Glycine-HCl and 1% (w/v) SDS, pH 3.0, and reprobed with anti-S6, anti-4E-BP, anti-STAT3 and anti-actin antibodies. Membranes were then processed as described above. Optical density of the relevant band was quantified using ImageJ 1.44c software (NIH).

### cDNA synthesis and Quantitative real-time PCR

Total RNA extracted from liver samples were treated with DNase I. Synthesis of cDNA was performed using the AMV reverse transcriptase according to the manufacturer’s instructions. The resulting cDNA was used for quantitative real-time PCR, using SYBR Green PCR Master mix. Thermal cycling was performed on QuantStudio 5 real-time PCR System (Thermo Fisher Scientific Inc.) using a relative calibration curve. The quantity of each mRNA transcript was measured and expressed relative to Glyceraldehyde-3-Phosphate dehydrogenase (GAPDH). The following primers used: Timp1: upstream 5′-GGT GTG CAC AGT GTT TCC CTG TTT-3′, downstream 5′-TCC GTC CAC AAA CAG TGA GTG TCA −3′^44^; Col4a1: upstream 5-CCA TGG TCA GGA CTT GGG TA −3′, downstream 5′-AAG GGC ATG GTG CTG AAC T-3′^45^; GAPDH: upstream 5′-CGA CTT CAA CAG CAA CTC CCA CTC TTC C-3′, downstream 5′-TGG GTG GTC CAG GGT TTC TTA CTC CTT-3′^46^.

### Preparation of solutions

Alcohol solution was prepared from absolute anhydrous alcohol (190 proof) diluted to 20% alcohol (v/v) in tap water. RapaLink-1 (1mg/kg)^21^, and RapaBlock (40mg/kg)^31^ were dissolved in 5% DMSO, 5% Tween 80, 5% PEG300 and 85% saline. Vehicle was consisted of 5% DMSO, 5% Tween 80, 5% PEG300 and 85% saline.

### Glucose tolerance test

Glucose tolerance test was performed as described previously^47^. Briefly, mice were deprived of food for 6 hours and then injected intraperitoneally (i.p.) with 1 g/kg glucose. Blood samples were taken from a tail vein nick at different time intervals (0, 15min, 30min, 60 min and 120min post glucose administration), and blood glucose level was analyzed using a Bayer Contour blood glucose meter and test strips.

### Behavioral testing

Mice were systemically administered with vehicle or RapaBlock (40 mg/kg) on Mondays, Wednesdays, and Fridays and tested after 3 weeks and 6 weeks of chronic drug treatment.

#### Rotarod test

The rotarod test was conducted as described previously^48^. Specifically, sensorimotor performance was assessed by the accelerating Rotarod apparatus (Rota-rod 7650, Jones & Roberts). Each trial started at 4 rpm and reached 40 rpm speed after 300 seconds. Mice underwent three trials, with 5-mininutes rest time in between trails. The trial ended when the mouse fell off the rod, completed one full revolution on the rod or when the speed of the apparatus reached 40 rpm (300 seconds). Latency to fall was scored in seconds, with 300 seconds as the maximum value.

#### Novel object recognition (NOR) test

The paradigm was conducted as described previously^49^, with small modifications. Mice were first acclimated to the experimental room for 60 minutes. Afterwards, two identical objects were placed in an open field (5 cm away from the walls), and mice were allowed to familiarize with both objects until they reached the criteria of 20 seconds of total exploration time. Six hours after the familiarization session, one familiar object was replaced by a novel object (the position of the novel object, left or right, was randomized between mice and groups). The mice were allowed to freely explore the open field until reaching the 20 seconds criterion of total exploration time. Exploration was characterized by the nose of the mouse directed toward an object at less than 2 cm of distance. Exploration time was recorded using the Ethovision XT video-tracking system (Noldus, Leesburg, VA, USA). The open field and the objects were cleaned with 75% ethanol and dried between each mouse and session.

#### Elevated plus maze (EPM) paradigm

The EPM paradigm was conducted as described previously^48^. Specifically, the EPM apparatus consists of two open and two closed arms (30 × 5 centimeters) with walls of 15 centimeters high and is elevated 40 centimeters above the ground. The arms extend from a central platform (5 × 5 centimeters) forming a plus sign. EPM testing took place in a quiet, dimly illuminated room. Each mouse was tested for 5 minutes after being placed in the center platform facing an open arm. The time spent on the closed arms and open arms of the EPM was scored. Arm entries were scored when an animal put all four paws into the arm. At the end of the test, the time spent in the open arms were expressed in seconds, and total distance traveled in centimeters. Data was recorder via Ethovision XT video-tracking system (Noldus, Leesburg, VA, USA). EPM apparatus was cleaned with 75% ethanol and dried between animals and sessions.

#### Alcohol Drinking paradigm

Mice underwent 7 weeks of intermittent access to 20% (v/v) alcohol in a 2-bottle choice drinking paradigm (IA20%2BC) as described previously^50^. Specifically, mice had 24-hour access to one bottle of 20% alcohol and one bottle of water on Mondays, Wednesdays and Fridays, with alcohol drinking sessions starting 2 hours into the dark cycle. During the 24 or 48 hours (weekend) of alcohol deprivation periods, mice had access to a bottle of water. The placement (right or left) of the bottles was alternated in each session to control for side preference. Two bottles containing water and alcohol in an empty case were used to evaluate the spillage. Alcohol and water intake were measured at the 4-hour and 24-hour time points. Alcohol preference ratio was calculated as the volume of alcohol intake/total volume of fluid intake (water+alcohol).

#### Drug administration

On week 8 following IA20%2BC, mice were systemically administered with vehicle (5% DMSO, 5% Tween 80, 5% PEG300 and 85% saline), RapaLink-1 (1 mg/kg), RapaBlock (40 mg/kg) or a combination of RapaLink-1 (1 mg/kg) and RapaBlock (40 mg/kg) 3 hours prior to the beginning of the drinking session, and alcohol and water intake were measured.

### Blood Alcohol Concentration Measurement

Blood alcohol concentration (BAC) procedure was conducted as described in^20^ with modifications. C57BL/6 mice were systemically co-administered with RapaLink-1 (1 mg/kg) and RapaBlock (40 mg/kg) or vehicle. Three hours after drug administration, mice received an i.p. injection of 2 g/kg of alcohol and blood was collected intracardially in heparinized capillary tubes 30 minutes later. Serum was extracted with 3.4% trichloroacetic acid followed by a 5-minute centrifugation at 420 g and assayed for alcohol content using the NAD-NADH enzyme spectrophotometric method^51^. BAC was determined by using a standard calibration curve.

### Statistical analysis

GraphPad Prism 7.0 (GraphPad Software Inc., La Jolla, CA) was used to plot and analyze the data. D’Agostino–Pearson normality test and F-test/Levene tests were used to verify the normal distribution of variables and the homogeneity of variance, respectively. Data were analyzed using the appropriate statistical test, including two-tailed unpaired t-test, one-way analysis of variance (ANOVA) and two-way ANOVA followed by post hoc tests as detailed in the Figure Legends. All data are expressed as mean ± SEM, and statistical significance was set at p<0.05.

---

**Fig. S1.**
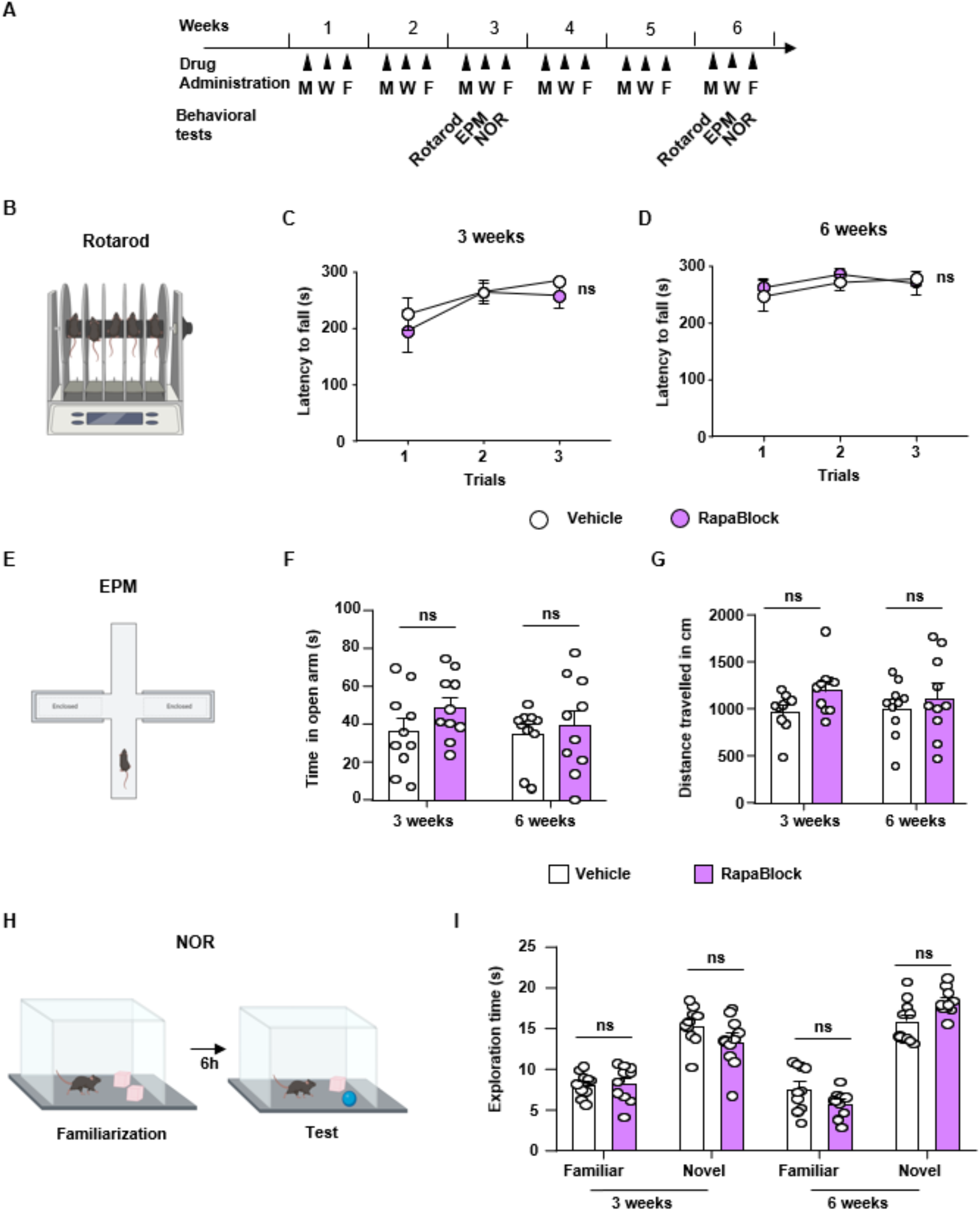
Chronic administration of RapaBlock has no adverse behavioral effects. **(A)** Timeline of experiments. Mice were treated with RapaBlock (40mg/kg, pink) 3 times a week for 6 weeks. Rotarod test **(B)**, elevated plus maze test (EPM) **(E)** and Novel object recognition test (NOR) test **(H)** were performed after 3 weeks and 6 weeks of treatment. (**C,D**) Performance on the accelerating Rotarod was not affected by chronic RapaBlock treatment (Two-way ANOVA, no effect of treatment after 3 weeks (*F*_1,18_ = 0.6013, *P* = 0.4482) and after 6 weeks (*F*_1,18_ = 0.1274, *P* = 0.7253)). **(F, G)**Time spent in the open arm of the EPM (RM Two-way ANOVA, (*F*_1,18_ = 2.435, *P* = 0.1361)) and total distance travelled during the test (RM Two-way ANOVA, (*F*_1,18_ = 4.183, *P* = 0.0557)) were not affected by chronic RapaBlock at both time points. **(I)** Time exploring the familiar and novel object were not affected by chronic RapaBlock treatment at both time points (RM Two-way ANOVA, (*F*_1,18_ = 0.7542, *P* = 0.3966)). Data are presented as individual data points and mean ± SEM, n=10 per condition. Significance was determined using RM Two-way ANOVA followed by Tukey’s multiple comparisons test. ns = non-significant.

**Fig. S2.**
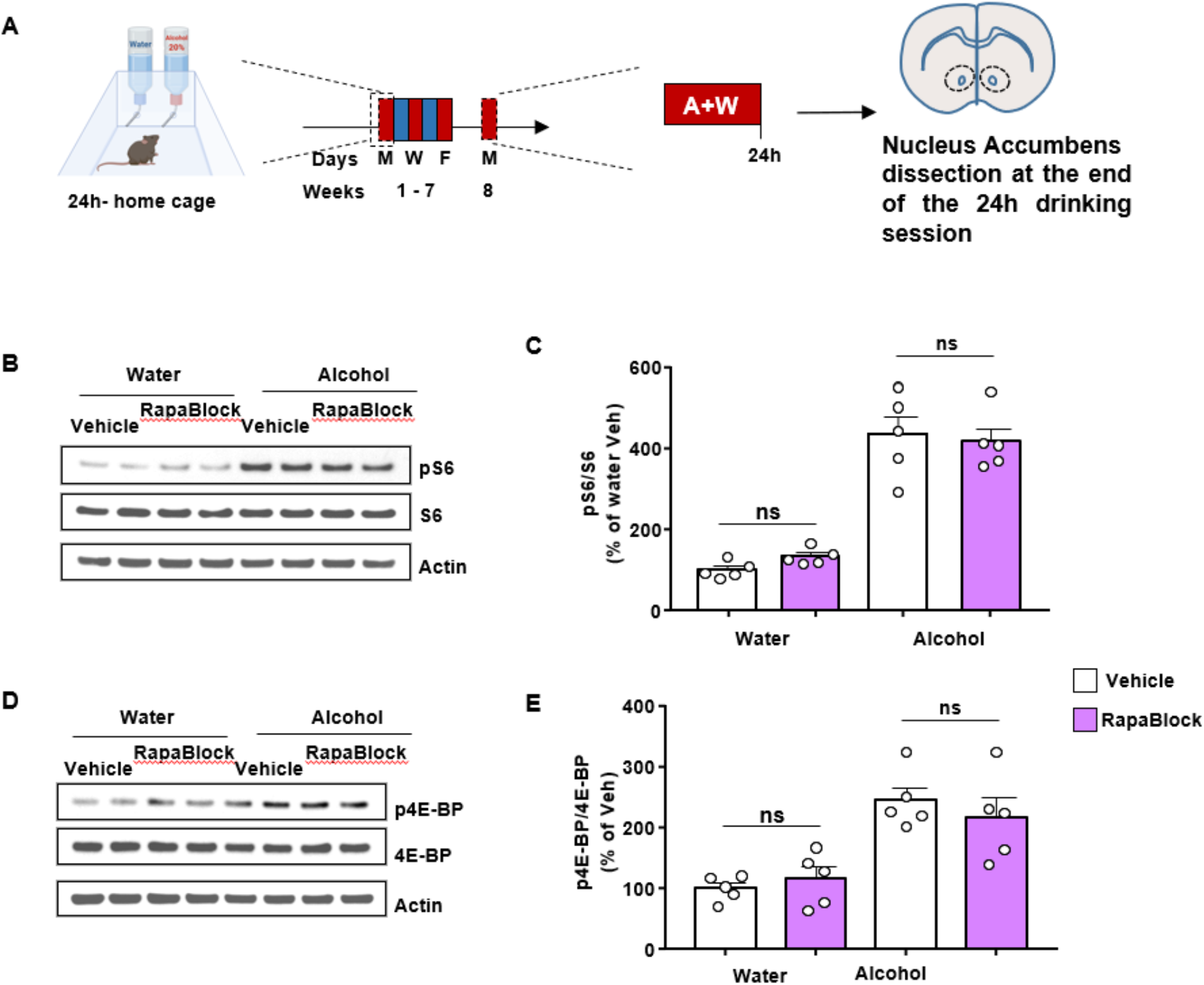
RapaBlock does not alter alcohol-dependent activation of mTORC1 in the Nucleus Accumbens. **(A)** Timeline of experiment. Mice underwent 7 weeks of IA20%2BC. On week 8, mice received a systemic administration of RapaBlock (40mg/kg) 3 hours before the beginning the session, and the NAc was removed at the end of the 24 hours drinking session. **(B,D)** Representative images of phospho-S6 (pS6) (**B**) and phospho-4E-BP (p-4E-BP) (**D**) (top panels), total protein levels of S6 (**B**) and 4E-BP (**D**) (middle panels), and actin (bottom panels). **(C,E)** Data are presented as the individual data points and mean densitometry values of the phosphorylated protein divided by the densitometry values of the total protein ± SEM and expressed as % of vehicle. Significance was determined using Two-way ANOVA. Administration of RapaBlock does not affect the levels of pS6 (Two-way ANOVA, main effect of alcohol (F(1,16)=270.2, p<0.0001), no effect of treatment (*F*_1,16_ = 0.1542, *P* = 0.6997)) and p4E-BP (Two-way ANOVA, main effect of alcohol (*F*_1,16_ = 37.31, *P* < 0.0001), no effect of treatment (*F*_1,16_ = 0.1707, *P* = 0.6849)). n=5 per condition. ns = non-significant.

**Fig. S3.**
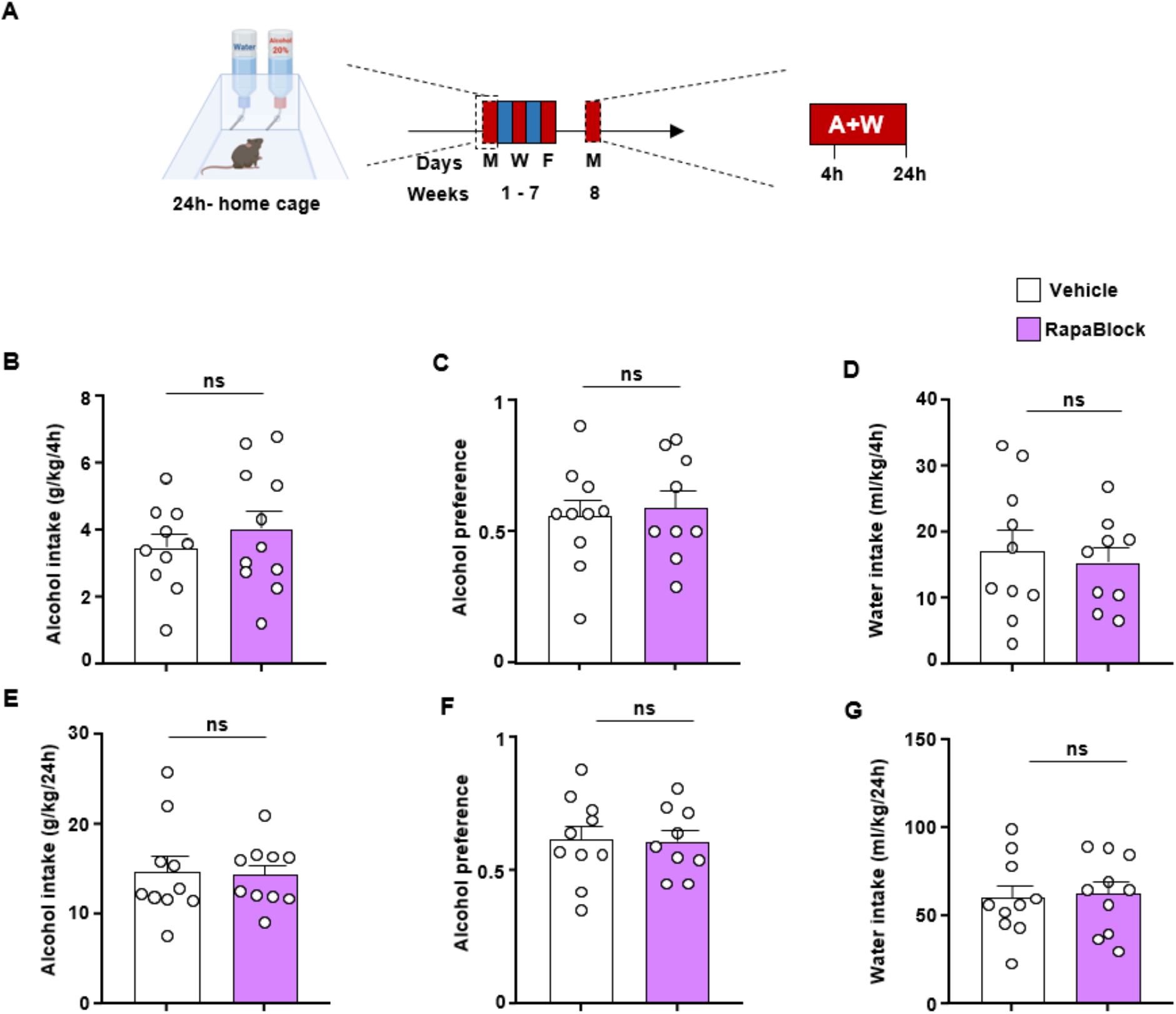
RapaBlock does not alter alcohol intake. **(A)** Mice underwent 7 weeks of IA20%2BC. On week 8, mice received a systemic administration of RapaBlock (40mg/kg) 3 hours before the beginning of drinking session. Alcohol and water intake were measured 4 hours **(B, D)** and 24 hours later **(E, G)**. Alcohol preference was calculated as the ratio of alcohol intake to total fluid intake at the 4-hour **(C)** and 24-hour **(F)** time points. **(B-D)** Administration of RapaBlock does not alter alcohol intake (two-tailed unpaired t-test: t = 0.4256, *P* = 0.6754), alcohol preference (Two tailed unpaired t-test: t = 0.3636, *P* = 0.7207) and water intake was (Two-tailed unpaired t-test: t = 0.4192, *P* = 0.6803) at the end of the 4-hour session. (**E-G**) Data are presented as individual data points and mean ± SEM. Significance was determined using Two-tailed unpaired t-test. Administration of RapaBlock does not alter alcohol intake (Two-tailed unpaired t-test: t = 0.1546, *P* = 0.8788), alcohol preference (Two-tailed unpaired t-test: t = 0.1191, *P* = 0.9066) and water intake (Two-tailed unpaired t-test: t = 0.2297, *P* = 0.8209) at the end of the 24-hour session. **(B-G)** n=9-10 per condition. ns = non-significant.

**Fig. S4.**
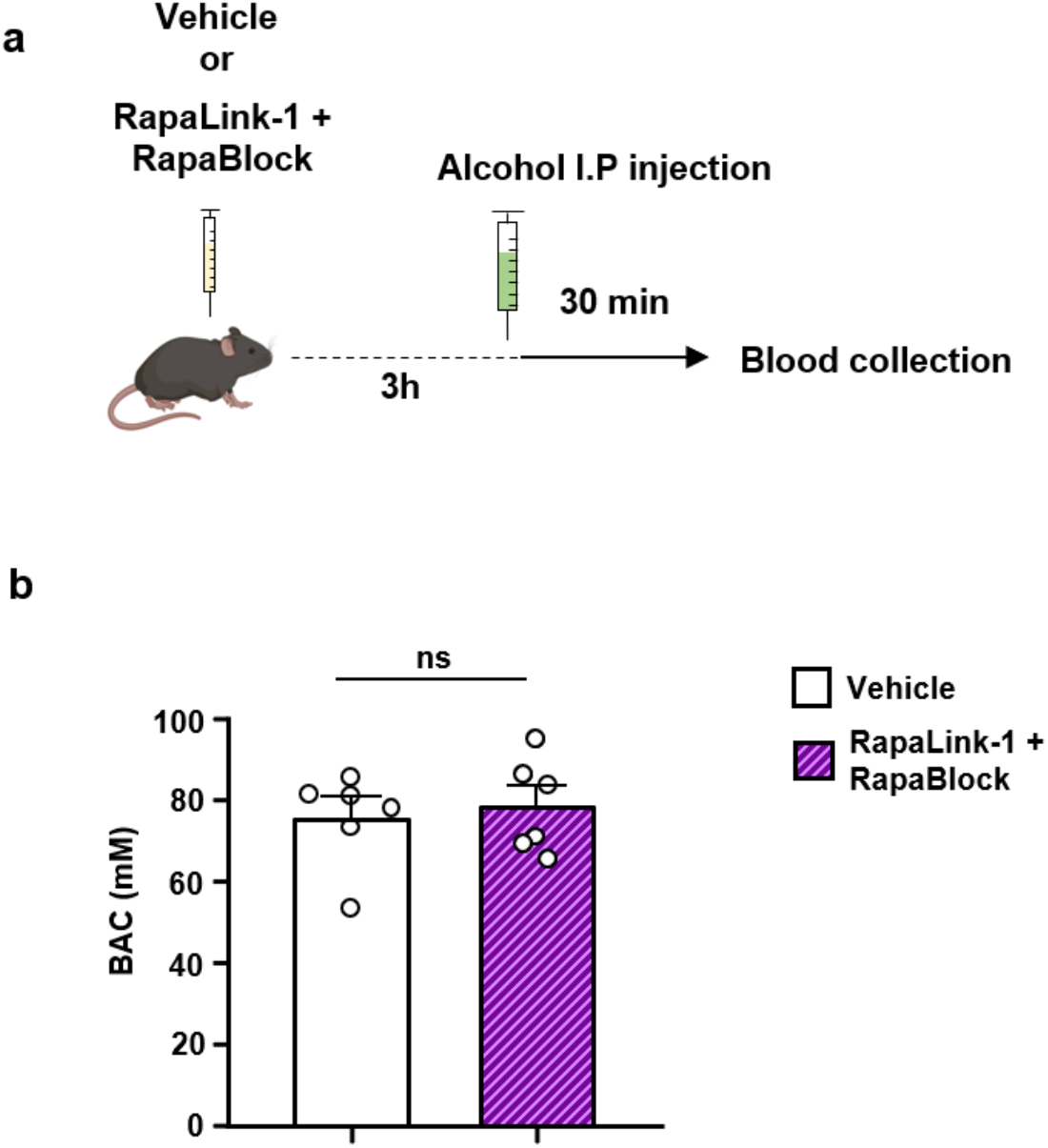
Systemic administration of a combination of RapaLink-1 and RapaBlock does not affect blood alcohol concentration. **(A)** Timeline of experiment. Mice received a systemic administration of vehicle or RapaLink-1 (1mg/kg, purple) + RapaBlock (40 mg/kg, pink). Three hour later, mice received a systemic administration of alcohol (2 g/kg), and BAC was measured 30 minutes later. **(B)** Data are presented as mean ± SEM. Significance was determined using Two-tailed unpaired t-test. BAC is similar in vehicle-treated vs. RapaLink-1 + RapaBlock-treated mice (Two-tailed unpaired t-test: t=0.4580, *P* = 0.6567). n = 6 per group. ns = non-significant.

